# Disequilibrium between chloroplast proton motive force and ATP levels in Arabidopsis

**DOI:** 10.1101/2025.11.12.687978

**Authors:** Gustaf E. Degen, Markus Schwarzländer, Matthew P. Johnson

## Abstract

Current dogma holds that CO_2_ fixation by photosynthesis requires additional ATP production via PGR5-dependent cyclic electron transfer (PGR5-CET) to augment the NADPH and ATP produced by linear electron transfer (LET). Here we use the MgATP^2−^ fluorescent biosensor ATeam1.03-nD/nA and find that while proton motive force (pmf) in the Arabidopsis *pgr5CAS* mutant is only 50-75% of that of the wild-type, MgATP^2−^ levels are unchanged. These data demonstrate that disequilibrium exists between pmf and MgATP^2−^ concentration in vivo, indicating that PGR5-CET is not required for ATP augmentation.

## Main text

CO_2_-fixation in plants via the Calvin-Benson-Bassham (CBB) cycle is fuelled by conversion of light energy into chemical energy and reducing power in the form of ATP and NADPH. NADPH is formed via linear electron transfer (LET) from water to NADP^+^ and is coupled to movement of 6H^+^ across the thylakoid membrane via the water-splitting reaction at photosystem II (PSII) and Q-cycle at cytochrome *b*_6_*f* (cyt *b*_6_*f*). The transmembrane proton motive force (pmf) is harnessed by the chloroplast ATP synthase to generate ATP from ADP and phosphate (Pi). Since the ATP synthase possesses 14 proton-binding c-subunits, the H^+^/ATP stoichiometry has been generally assumed to be 4.7 (14 divided by the 3 ATP molecules formed per 360° rotation of the enzyme)^1^. Therefore, 1.28 ATP (6/4.7) can be formed per 1 NADPH ^2^. However, the CBB cycle requires 9 ATP and 6 NADPH for CO_2_ fixation—a ratio of 1.5^3^. This is the central bioenergetic dogma underpinning oxygenic photosynthesis.

Any shortfall in ATP synthesis relative to NADPH production requires either the generation of additional pmf or the export of reductant from the chloroplast. In C3 angiosperms additional pmf can be generated via cyclic electron transfer (CET)^4^. CET involves the movement of electrons from the photosystem I (PSI) electron acceptor ferredoxin (Fd), which normally reduces NADP^+^, instead to the cyt *b*_6_*f* electron donor plastoquinone (PQ). This ensures that electrons pass through the Q-cycle a second time, thus increasing the H^+^/e^−^ stoichiometry. In C3 angiosperms, CET is mediated mainly by PROTON GRADIENT REGULATION 5 (PGR5) and to a lesser extent by NADH dehydrogenase-like complex (NDH)^5^. *Arabidopsis thaliana* mutants lacking PGR5 have been generated by ethyl methane sulfonate *(pgr5)* and CRISPR/CAS9-engineering (*pgr5CAS)*, and these plants show lower pmf, LET, CO_2_-fixation rates, and over-reduction of PSI electron acceptors consistent with a shortfall in ATP augmentation due to loss of CET^6,7^. However, whilst these phenotypes have been measured using well-established gas-exchange, chlorophyll fluorescence and absorption techniques, direct evidence of a shortfall in ATP production in *pgr5* and *pgr5CAS* plants has never been provided *in vivo*.

An interesting and consistently observed feature of *pgr5* and *pgr5CAS* plants is their much higher proton conductivity (gH^+^) compared to the wild-type at moderate to high light intensities^8^. High gH^+^ could indicate a lower phosphorylation potential in *pgr5* chloroplasts (less displacement of the ATP→ADP+Pi reaction from equilibrium) due to lower ATP concentration^8^. Indeed, it has been suggested that ATP synthase in *pgr5* is ‘leaky’ (i.e. exhibits a non-productive proton leak), but this hypothesis was later ruled out through analysis of double mutants^9^. Alternatively, it was suggested that PGR5 might negatively regulate ATP synthase, restricting gH^+^ to allow pmf build-up, which would imply higher ATP levels in the *pgr5* mutant (a productive proton ‘leak’)^10^.

Since CET generates additional pmf, of which the major component is the proton concentration gradient (ΔpH) in chloroplasts, it also plays a key role in triggering ΔpH-dependent photoprotective mechanisms. These include photosynthetic control (PCON) which controls the rate of electron transfer in the cyt *b*_6_*f* complex, preventing overreduction and damage to PSI^11^; and nonphotochemical quenching (NPQ), which protects PSII by dissipating excess excitation energy with its light harvesting antenna system^12^. Consistent with this concept, *pgr5* and *pgr5CAS* plants suffer from photoinhibition of photosystem I (PSI) and a strong decrease in the level of NPQ^5,13^. Therefore, loss of the additional pmf required for triggering photoprotective mechanisms on its own may explain the low LET and CO_2_ fixation phenotype in both *pgr5* mutants^14–16^.

Clearly new approaches are required to distinguish between whether PGR5 plays a key role in ATP augmentation, in photoprotection or both. Recently, a MgATP^2−^ fluorescent biosensor (ATeam1.03-nD/nA) was successfully expressed in Arabidopsis chloroplasts via the transketolase chloroplast transit peptide (TKTP) demonstrating how MgATP^2−^ levels change upon illumination due to LET activity ^17,18^. The MgATP^2−^ biosensor relies on FRET transfer between a monomeric super-enhanced cyan fluorescent protein (CFP) and circularly permuted monomeric Venus (a yellow fluorescent protein variant, YFP), which are both fused to an MgATP^2−^ sensing epsilon subunit of a bacterial ATP synthase^18^. Binding of ATP will bring both fluorophores into closer proximity increasing FRET efficiency, i.e. upon excitation of CFP at 458 nm energy will be transferred from CFP to YFP, resulting in an increased YFP/CFP emission ratio with increasing MgATP^2−^ concentration. Here we establish the ATeam1.03-nD/nA fluorescent biosensor as a new tool in photosynthesis research to understand how MgATP^2−^ levels are correlated with pmf, gH^+^ and proton flux (vH^+^) across the thylakoid membrane in WT and *pgr5CAS* plants.

In Figure 1a we show confocal fluorescence microscopy images of the TKTP-ATeam1.03-nD/nA biosensor in the wild-type background (hereafter WT-TA), as previously reported the CFP and YFP fluorescence signals are co-located with the chlorophyll fluorescence confirming their localisation in the chloroplast ^18^. We transformed the *pgr5CAS* mutant with the same construct to obtain two independent lines, *pgr5CAS* TKTP-ATeam 25 and 1513 (hereafter *pgr5CAS*-TA1 and *pgr5CAS*-TA2, respectively) (Figs 1a, S2a). As with the WT, the YFP and CFP fluorescence signals co-locate with the chlorophyll autofluorescence of the chloroplast. We next performed growth assays to confirm that the transformed WT and *pgr5CAS*-TA1 and *pgr5CAS*-TA2 plants did not show perturbed growth compared to the parent background strains (Figs 1b, S2b).

**Figure 1.**
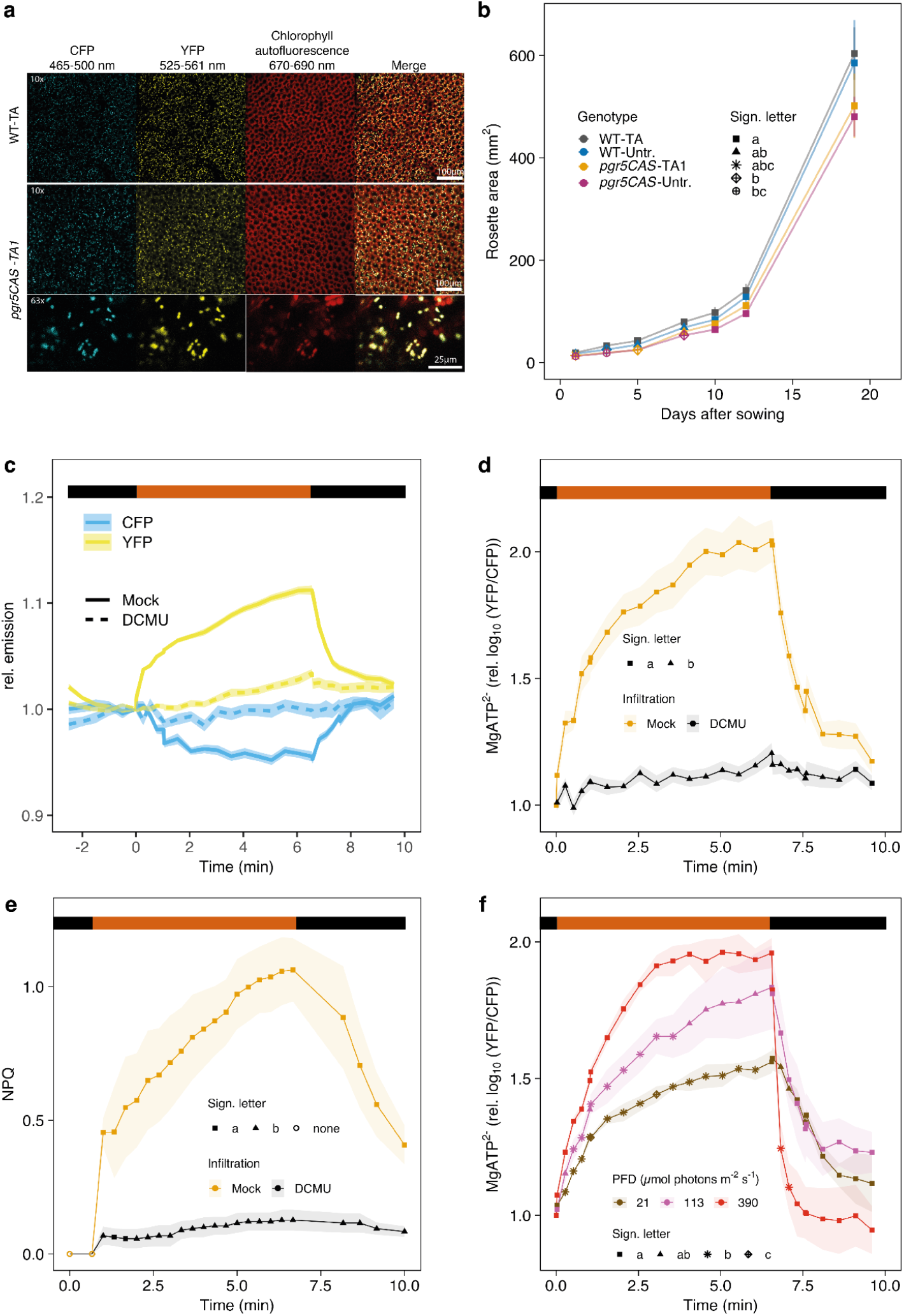
Light-dependent MgATP^2−^ accumulation in *pgr5CAS*-TA1. a) Confocal images of WT-TA and *pgr5CAS*-TA1 lines transformed with the chloroplast stroma-localised FRET MgATP^2−^ biosensor ATeam1.03-nD/nA. b) Rosette area of untransformed WT and *pgr5CAS* lines and transformed lines shown in panel (a). c) Intensities of the single CFP and YFP channels during dark-light-dark transitions, normalised to the last dark time point (t=0) of *pgr5CAS*-TA1 of leaves infiltrated with Mock or 20 µM DCMU. d) MgATP^2−^ dynamics derived from normalised log_10_-transformed FRET ratios of Mock and DCMU-treated *pgr5CAS-TA1* leaves during dark-light-dark transitions. e) Nonphotochemical fluorescence quenching of leaves from the same plants and treatments as in (c). f) MgATP^2−^ dynamics of *pgr5CAS-TA1* leaves during three light intensities. Data points represent the mean of at least 3 biological replicates and shaded areas represent SEM. Different symbols indicate significant differences between treatments or light intensities at each time point calculated from a Tukey HSD test, alpha = 0.05. For (c)-(d) the red bar corresponds to 390 µmol photons m^−2^ s^−1^ of red actinic light, for (f) the red bar indicates when the light was turned on.

Since confocal fluorescence microscopy measurements required detached leaves sandwiched between glass coverslips (limiting air contact; Fig S1a), we sought to identify conditions yielding results comparable to those from attached leaves. We found that incubating leaves with 100 mM NaHCO_3_ (as a CO_2_ source) yielded results similar to those from attached leaves (Fig S1b); therefore, we used this concentration in all subsequent experiments. Previous work showed that the TKTP-ATeam1.03-nD/nA biosensor affinity (K_d_) for MgATP^2−^ decreases with increasing temperature; therefore, we rigorously controlled temperature, ensuring it did not exceed 22.5 °C even at the highest light intensity (Fig S1c). We then validated that the ATeam biosensor had sufficient dynamic range by infiltrating leaves with 0, 0.8, 1.6 and 3.2 mM equimolar ATP and MgCl_2_ in the presence of digitonin (Fig S1d), confirming that the sensor is well within the range of previously reported stromal ATP concentration in the light of 0.5-1.8 mM^19–22^.

We next confirmed that the ATeam1.03-nD/nA biosensor responded to light in the *pgr5CAS-* TA1 mutant (Fig 1c). As expected, the CFP fluorescence decreased and the YFP fluorescence increased during a dark to light transition and then returned to close to the dark baseline upon cessation of illumination (Fig 1c). This change was sensitive to 20 μM of the herbicide *n*-(3,4-dichlorophenyl)-*n*-dimethylurea (DCMU) which poisons PSII and thus blocks LET (Fig 1c). The ratio of the YFP/CFP fluorescence indicative of the MgATP^2−^ level in the chloroplast thus increased in the light and decreased again upon return to darkness (Fig 1d) and both this change and the accompanying change in NPQ were eliminated by DCMU (Fig 1e). The MgATP^2−^ level in the *pgr5CAS*-TA1 mutant also showed a response which scaled with increasing light intensity consistent with increased proton-coupled photosynthetic electron transfer (Fig 1f). MgATP^2−^ levels decline more rapidly in the dark following higher intensity illumination, consistent with enhanced CO_2_ fixation rate resulting from increased CBB cycle activation.

Subsequently, we confirmed the photosynthetic phenotype of the *pgr5CAS*-TA1 plants was consistent with the previously reported phenotype for the *pgr5CAS* mutant using chlorophyll fluorescence, P700 and electrochromic shift (ECS) absorption measurements at light intensities of 0, 54, 83, 178, 228, 390 µmol photons m^−2^ s^−1^. Compared to the WT-TA, the *pgr5CAS*-TA1 plants showed decreased NPQ (Fig 2a) and lower PSII quantum yield (YII) (Fig S3a) with increasing light intensity. Similarly, *pgr5CAS*-TA1 showed faster P700 oxidation under far-red light indicative of lower CET (Fig S3b), and under red light lower PSI quantum yield (YI) (Fig S3c), decreased PSI donor side limitation (YND) (Fig S3d) and increased PSI acceptor side limitation (YNA) (Fig S3e). Consistent with this phenotype, the *pgr5CAS*-TA1 plants showed lower pmf with increasing light intensity compared to WT-TA (Fig 2b), mostly because of lower proton flux (vH^+^) consistent with a loss in CET (Fig S3f).

**Figure 2.**
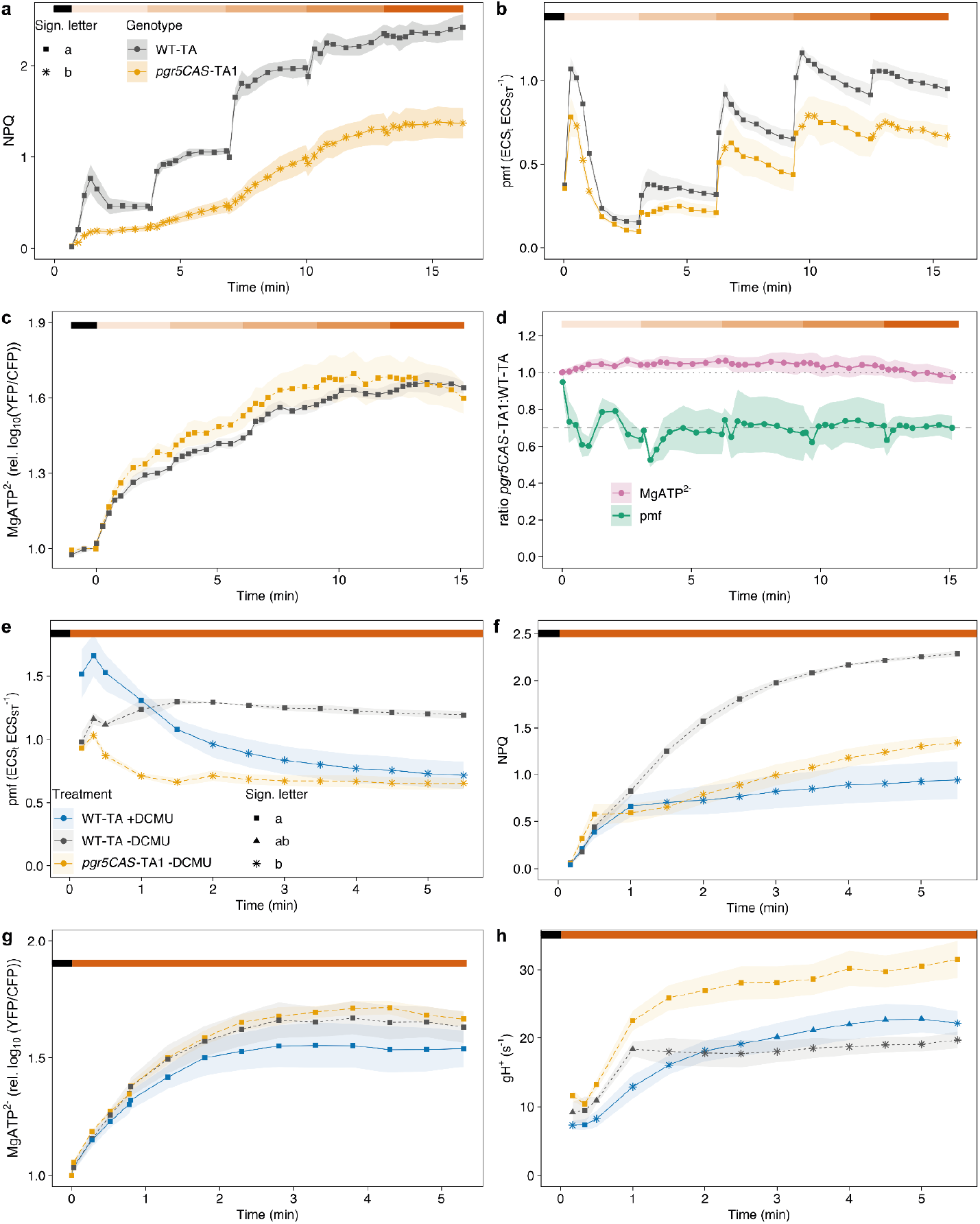
Photosynthetic parameters and MgATP^2−^ dynamics in WT-TA and *pgr5CAS*-TA1 plants. a) Nonphotochemical fluorescence quenching (NPQ). b) Proton motive force (pmf) calculated as ECS_t_ ECS_ST_^−1^. c) MgATP^2−^ dynamics derived from the chloroplast stroma-localised FRET MgATP^2−^ biosensor ATeam1.03-nD/nA. Log_10_-transformed FRET-ratios relative to the last dark time point (t=0) prior to onset of illumination are shown. d) Ratio of *pgr5CAS*-TA1 MgATP^2−^ and pmf levels relative to WT TKTP-ATeam at increasing light intensities derived from panels (b) and (c). e) pmf during dark-to-light transitions of WT TKTP-ATeam and *pgr5CAS-TA1* infiltrated without or with 4 µM DCMU. f) NPQ of the same plants and treatments as in (e). g) MgATP^2−^ dynamics of the same plants and treatments as in (e). h) Proton conductance of the thylakoid membrane of the same plants and treatments as in (e). Data points represent the mean of at least 3 biological replicates and shaded areas represent SEM. Different symbols indicate significant differences between treatments or genotypes at each time point calculated from a Tukey HSD test, alpha = 0.05. Coloured bars at the top of the panel indicate red actinic light intensities. For (a) (Dual KLAS, peak *λ* = 622): 0, 59, 87, 163, 294, 383 µmol photons m^−2^ s^−1^. For (b) (Dual PAM, peak *λ* = 622): 0, 50, 77, 152, 287, 378 µmol photons m^−2^ s^−1^. For (c) (LED ring combined with confocal microscope, peak *λ* = 644): 0,54, 83, 178, 228, 390 µmol photons m^−2^ s^−1^. The light intensities during induction shown in (e-h) were 383, 378, 390 and 378 µmol photons m^−2^ s^−1^, respectively.

Despite these differences in key photosynthetic parameters and pmf, we found the *pgr5CAS*-TA1 plants showed very similar chloroplast MgATP^2−^ levels compared to the WT-TA (Fig 2c). In both cases the increasing light intensity led to an increase in MgATP^2−^ levels with saturation reached at 228 µmol photons m^−2^ s^−1^ for both sets of plants. The disparity between pmf and MgATP^2−^ levels is shown in Fig 2d, where *pgr5CAS*-TA1 plants maintain ∼100% of the WT ATP level with only ∼70-75% of the pmf.

Our biosensor data reveal significant disequilibrium between thylakoid pmf and chloroplast MgATP^2−^ levels *in vivo*. We tested this further by infiltrating WT-TA leaves with subsaturating 4 μM DCMU. Partial inhibition of photosynthetic electron transfer lowered pmf in the WT by ∼50%—to a level consistent with that observed in *pgr5CAS*-TA1 after 5 min illumination at 390 µmol photons m^−2^ s^−1^ (Fig 2e). The lower pmf in the 4 μM DCMU treated WT-TA leaves also lowered NPQ to a level commensurate with that in *pgr5CAS*-TA1 (Fig 2f). Nonetheless, despite the 50% drop in pmf, ATP levels in the 4 μM DCMU-treated WT-TA leaves were not significantly different from those in untreated WT-TA or *pgr5CAS*-TA1 (Fig 2g). Interestingly, despite a significantly higher gH^+^ in *pgr5CAS*-TA1 compared to the WT-TA the MgATP^2−^ levels remain similar (Fig 2h). In the independent *pgr5CAS*-TA2 line, we also observed a significantly decreased pmf and increased gH^+^ but no differences in MgATP^2−^ levels compared to WT-TA (Fig S2c-e).

Overall, our results overturn two decades of dogma in photosynthesis research that PGR5-CET is necessary to augment chloroplast ATP levels for maximal CO_2_ fixation. Instead, they reveal disequilibrium between thylakoid pmf and stromal ATP levels *in vivo*, with just 50% of the WT pmf level sufficient to support ∼100% of the maximum MgATP^2−^ level we were able to observe in our experimental set-up. These data are consistent with past *in vitro* work on intact chloroplasts using biochemical methods to determine ATP concentration and dye-based spectroscopy to determine ΔpH^23,24^. An important caveat is that the TKTP-ATeam biosensor measures chloroplast MgATP^2−^ levels rather than ATP/ADP ratio or adenylate charge. We cannot therefore rule out that the thylakoid pmf is in overall equilibrium with stromal adenylate charge through the activity of the chloroplast adenylate kinase (AK), which could buffer MgATP^2−^ levels by the reaction ATP+AMP⇄ ADP+ADP^25^. For instance, it is possible that in *pgr5* the effect of loss of pmf is buffered by downregulation of stromal AK activity compared to the WT, thus less ATP is converted to ADP. However, our recent proteomic study of *pgr5* found no change in chloroplast AK levels^7^. Moreover, since the AK reaction dissipates pmf (upregulation of ATP synthase gH^+^ via [ADP] increase) one would expect a decrease in gH^+^ in *pgr5*, where in fact an increase is consistently observed. Indeed, if the WT chloroplast AK reaction is significant, under our conditions this should lead to dissipation of excess pmf. Yet pmf is considerably higher in WT than in *pgr5CAS* (25–50% higher)—inconsistent with significant AK activity. Rather, our data are consistent with the view that MgATP^2−^ and adenylate charge in the chloroplast are closely correlated as previously found intact chloroplasts in a range of studies^24,26,27^.

Our findings reframe our understanding of the Arabidopsis *pgr5* phenotype, suggesting that the photosensitivity and impairment to CO_2_ fixation at high light intensities observed derives not from an ATP shortfall but rather from the loss of the additional ΔpH required to activate photoprotective processes (Figure S4). Indeed, this neatly explains why the *pgr5* phenotype is ameliorated by processes that boost ΔpH such a flavodiiron-protein dependent pseudocyclic electron transfer (FLV-PCET) ^28^. Our biosensor data suggest that ATP synthase activity is downregulated in the WT above a certain threshold of ATP concentration, consistent with reports of so-called ‘metabolic control’ of its proton conductivity ^29,30^. This metabolic control of ATP synthase acts to preserve pmf (specifically ΔpH) for the regulation of photosynthesis via PCON and NPQ.

Whether PGR5 plays a role in direct regulation of the ATP synthase activity or alternatively plays a part in CET remains to be clarified. Our data (Fig 2h) suggest that any putative regulation of ATP synthase activity by PGR5 does not result in a productive proton leak in its absence, since elevated gH^+^ in *pgr5CAS*-TA1 or *pgr5CAS*-TA2 was not accompanied by higher ATP levels. Another alternative is that the high gH^+^ seen in *pgr5* reflects a genuine non-productive proton leak, though this seems unlikely since it is not observed in the *pgr5* background when FLV-PCET is introduced^28^. Saliently, the recent analysis of another high gH^+^ Arabidopsis mutant, *hope2*, revealed that it greatly increased PGR5 levels to compensate^7^. While this could be interpreted as a futile effort to restore gH^+^ control, the analysis of *hope2* revealed an increased rate of CET that was dependent on the presence of PGR5 and was successful in maintaining WT levels of pmf and NPQ ^7,31^. Collectively, these data argue for a role of PGR5 in CET to provide the additional proton influx necessary to build pmf, while metabolic regulation of the ATP synthase controls proton efflux, the two regulatory systems working synergistically to ensure photoprotection. The source of the elevated gH^+^ in *pgr5* now requires further investigation. One possibility is that altered ECS decay kinetics give the appearance of a leak, e.g. via altered Q-cycle dependent charge separation resulting from mis-regulation of cytochrome *b*_6_*f* in the absence of PGR5 ^32^.

Recently, biosensor approaches monitoring chloroplast NADPH levels (TKTP-iNap) and cytosolic NADH/NAD^+^ showed that re-oxidation of chloroplast NADPH relies in part on export of reducing equivalents to the cytosol via the malate valve and subsequent oxidation in the mitochondria ^33^. These data and our conclusions here collectively argue for a model where the ATP/NADPH ratio requirements of the CBB cycle in Arabidopsis are balanced primarily by exporting reducing equivalents rather than via augmentation of ATP within the chloroplast.

In conclusion, we show that the primary role of PGR5 is not in ATP augmentation but rather in providing additional ΔpH, that is required to reach the threshold required to activate photoprotective PCON and NPQ.

## Methods

### Generation of *pgr5CAS* lines expressing TKTP-ATeam and plant growth

Seeds lacking the PGR5 protein^16^, (generated via CRISPR-CAS9 and referred to as *pgr5CAS*) were sterilised in 70% (v/v) EtOH and 0.1% (v/v) Triton-X 100 and stratified at 4°C for 48 h. Seeds were sowed on Levington Advance Seed & Modular F2S Compost + Sand and placed under long-day conditions (16h light/ 8h dark, 22°C) and grown until flowering. For floral dip, *Agrobacterium tumefaciens* GV3101 cells were transformed with pEarleyGate100 containing the ATeam1.03-nD/nA sequence fused to the leader sequence from *Nicotiana tabacum* transketolase for targeting the plastid stroma under the control of a 35S promoter. T1 seeds were selected via resistance to glufosinate ammonium (120 mg L^−1^, 0.05 % Silvet-77 (v/v)). Resistant plants with high TKTP-ATeam fluorescence were selected and backgrounds from two independent transformation events were taken to homozygosity and used for the analyses. The line *pgr5CAS-*TA1 exhibited the strongest fluorescence and displayed a low level of generational silencing. The second line *pgr5CAS*-TA2 was used for additional validation experiments (Fig S2). For confocal imaging and PAM measurements, *Arabidopsis thaliana* WT and *pgr5CAS* expressing the TKTP-ATeam biosensor were grown in Levington Advance Seed & Modular F2S Compost + Sand under short-day conditions (9h light/ 15 dark, 23°C/15°C) for 4-5 weeks. Detached leaves were placed between two cover slips in the presence of 100 mM NaHCO_3_ as a CO_2_-source during confocal imaging and PAM measurements (validation shown in Fig. S1d).

### Confocal imaging

All confocal imaging was performed with a Zeiss Airyscan LSM880 in combination with a custom-built LED illumination chamber (Fig. S1a). CFP was excited at 458nm and emission was collected at 465-500 nm (CFP), 525-561 nm (YFP) and 670-700 nm for chlorophyll fluorescence. The pinhole was adjusted to 150 µm, pixel time to 2.05 μs and time to scan a full frame to 1.5s. For FRET measurements, an EC Plan-Neofluar 10x/03 objective was used and for localisation of the TKTP-ATeam biosensor to the chloroplast a Plan-Apochromat 63x/1.4 Oil DIC M27 objective was used. Detached leaves were placed between two cover slips in the presence of 100 mM NaHCO_3_ to avoid CO_2_-limitation. This was placed in a custom-made LED illumination chamber, which was integrated with the trigger interface of the Zeiss Zen software as described ^34^ (Fig. S1a). Confocal images were processed with a MATLAB-based Redox Ratio Analysis software (RRA)^35^. First, the background signal was removed using the integrated functionality of the RRA software. Then, for each biological replicate, 12 regions of interest (corresponding to 12 individual chloroplasts) were selected and the intensities of the single YFP and CFP channels were exported and used for further analysis and plotting in RStudio. The YFP/CFP intensity ratio was log_10_-transformed to approach the normal distribution and normalised to the value at time 0, the last time point in the dark prior to onset of illumination. For DCMU measurements, detached leaves were incubated in the dark for 40 min in water containing 4 or 20 µM DCMU and 0.2% EtOH (v/v) or 0.2 % EtOH (v/v) as a mock control. Spectra of the LEDs of the custom-built LED rings and Walz machines (described below) were determined using a LI-180 (LI-COR, Lincoln, United States). To measure the impact of LED illumination on the temperature of leaves mounted between two cover slips, a thermocouple was placed between the cover slips onto the leaves and temperatures were measured at increasing light intensities after 3 min at each light intensity (Fig. S1c)

### Photosynthesis measurements

The ECS at 515 nm was measured using a Dual-PAM analyser with a P515/535 emitter/detector module (Heinz Walz GmbH, Effeltrich, Germany) ^36^. Plants were dark-adapted for at least 1 h prior to measurements, and leaves placed between two cover slips, as described above. Proton motive force (pmf) was calculated from the amplitude of a single exponential decay fit to the first 500 ms of the ECS decay in the dark (ECS_t_), normalised to the height of a 50 μs single turnover flash (ECS_ST_) applied prior to onset of actinic light. Conductance of the thylakoid membrane to protons (gH^+^) was calculated as the inverse of the rate constant τ_ECS_ of the single exponential decay and proton flux was calculated as vH^+^= pmf × gH^+^.

A Dual-KLAS-NIR photosynthesis analyser (Heinz Walz GmbH, Effeltrich, Germany) was used for pulse-amplitude modulation chlorophyll fluorescence measurements and P700 absorption spectroscopy in the near-infrared ^37^. Measurements were performed as described in ^7^. Briefly, after plants had dark-adapted for at least 1 h, detached leaves were placed in 100 mM NaHCO_3_ between two cover slips and four pairs of pulse-modulated NIR measuring beans were zeroed and calibrated before each measurement. For each genotype, one leaf was used to generate differential model plots according to manufacturer’s protocol, which were used for online deconvolution to determine redox changes of P700. Prior to each measurement, maximum oxidation of P700 was determined by using the preprogrammed NIRmax routine. DCMU treatments for photosynthesis measurements were performed as described above.

### Statistical analysis

Statistical analysis was performed in RStudio. The aov function was used in conjunction with the TukeyHSD function to calculate the Tukey Honest Significant Differences, with the confidence level set to 0.95. Then, the multcompView4 function from the multcompLetters package was used to convert significant differences into a letter summary. Common letters between groups (i.e. a, ab, b) indicate non-significance, whereas distinct letters (i.e. a, b, c) indicate significant differences between groups.

## Acknowledgements

The authors would like to thank Dr Ke Zheng (University of Munster, Germany) for his advice on illumination microscopy. MPJ acknowledges funding from the Leverhulme Trust grant RPG-2021-345 and Biotechnology and Biological Sciences Research Council (BBSRC) grant UKRI1945. MS is grateful for funding by the German Research Foundation (DFG) through the Research Unit FOR 5573 ‘Dynamic regulation of the proton motive force in photosynthesis’ (SCHW 1719/11-1; SCHW 1719/12-1).

---

**Figure S1.**
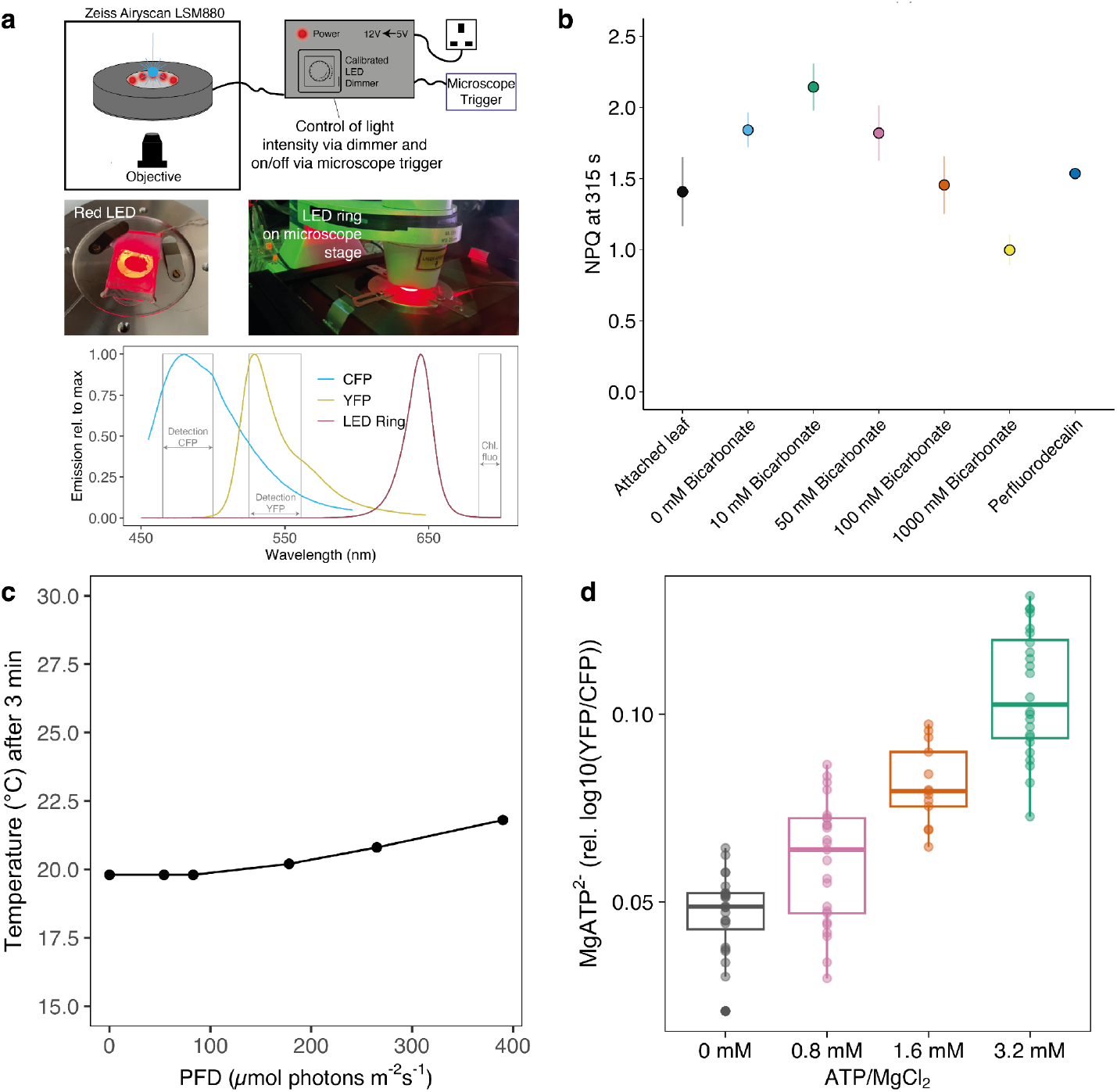
Experimental setup of the ATP biosensor. a) Top: Combination of the Zeiss Airyscan LSM880 with a red dimmable LED ring. The 12V LEDs integrate with the Zeiss ZenBlack software via the Trigger.Out interface. This allows switching the LEDs on/off at desired times. Light intensities can be manually controlled via the dimmer up to a maximum light intensity of 390 µmol photons m^−2^ s^−1^. Bottom: Emission spectrum of the biosensor and LED ring and detection windows. b) Validation of the experimental setup to include 100 mM NaHCO_3_ to avoid CO_2_-limitation of the leaf mounted between two cover slips. At 100 mM NaHCO_3_, similar NPQ values of WT-TA to an attached leaf were achieved. c) Temperature effect of the LED on the leaf mounted between two cover slips. The temperature of the leaf was determined after 3 min at each light intensity, corresponding to the setup used in Fig2. d) Response of the ATP biosensor in WT-TA leaves infiltrated with equimolar concentrations of ATP and MgCl_2_ in the presence of 100 µM digitonin to partially permeabilise the cell membranes and to test the responsiveness of the sensor to ATP increase.

**Figure S2.**
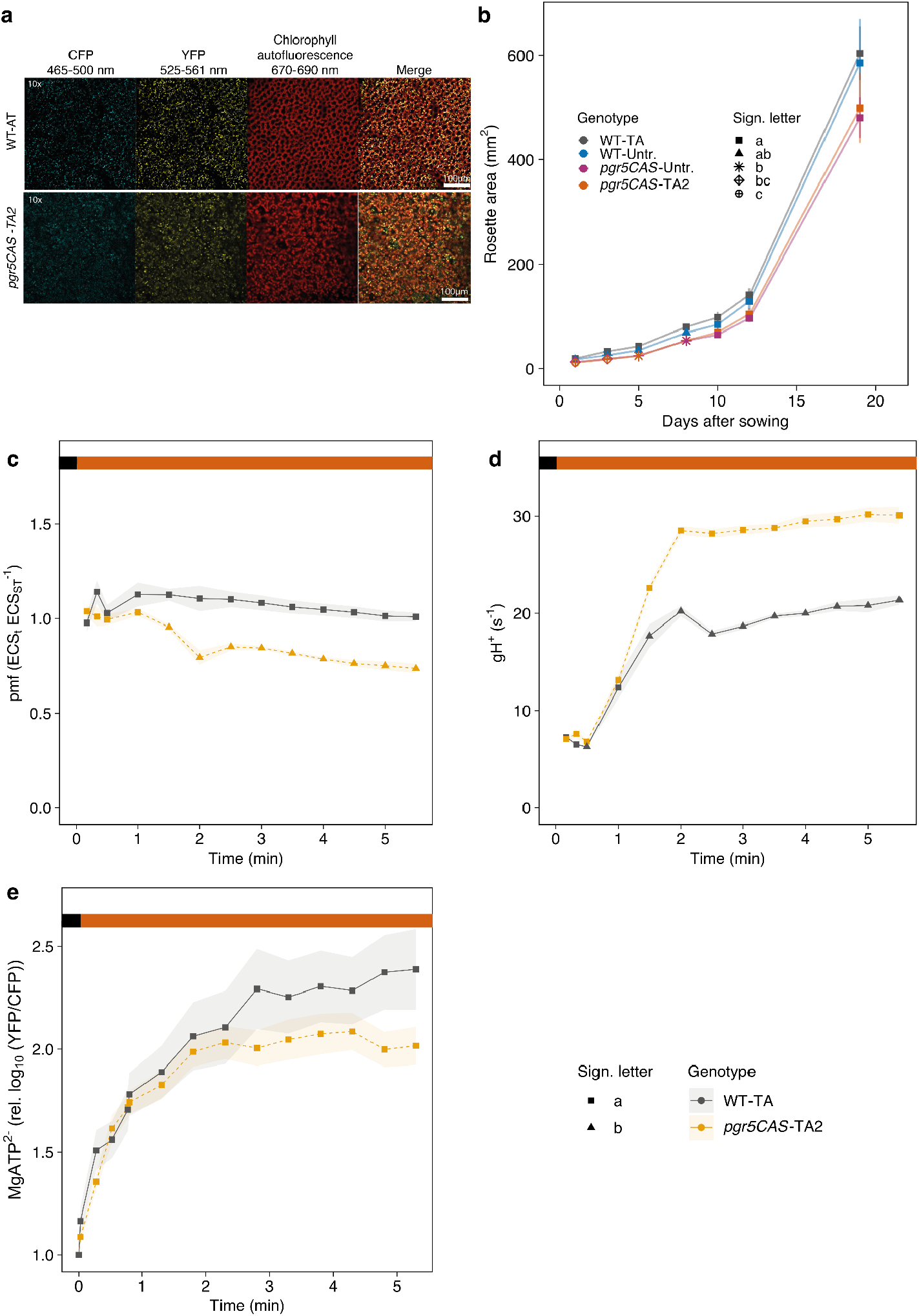
Validation of a second *pgr5CAS*-TA2. a) Confocal microscopy of WT-TA and *pgr5CAS*-TA2 showing the stable expression of the chloroplast stroma-localised FRET MgATP^2−^ biosensors ATeam1.03-nD/nA. b) Rosette area between untransformed and transformed WT and *pgr5CAS* lines. c) pmf in WT-TA and *pgr5CAS*-TA2, d) gH^+^ in WT-TA and *pgr5CAS*-TA2, e) MgATP^2−^ levels in WT-TA and *pgr5CAS*-TA2. Data points represent the mean of at least 3 biological replicates and shaded areas represent SEM. Different symbols indicate significant differences between treatments or genotypes at each time point calculated from a Tukey HSD test, alpha = 0.05. Coloured bars at the top of the panel indicate red actinic light intensities used in Fig 2e-h.

**Figure S3.**
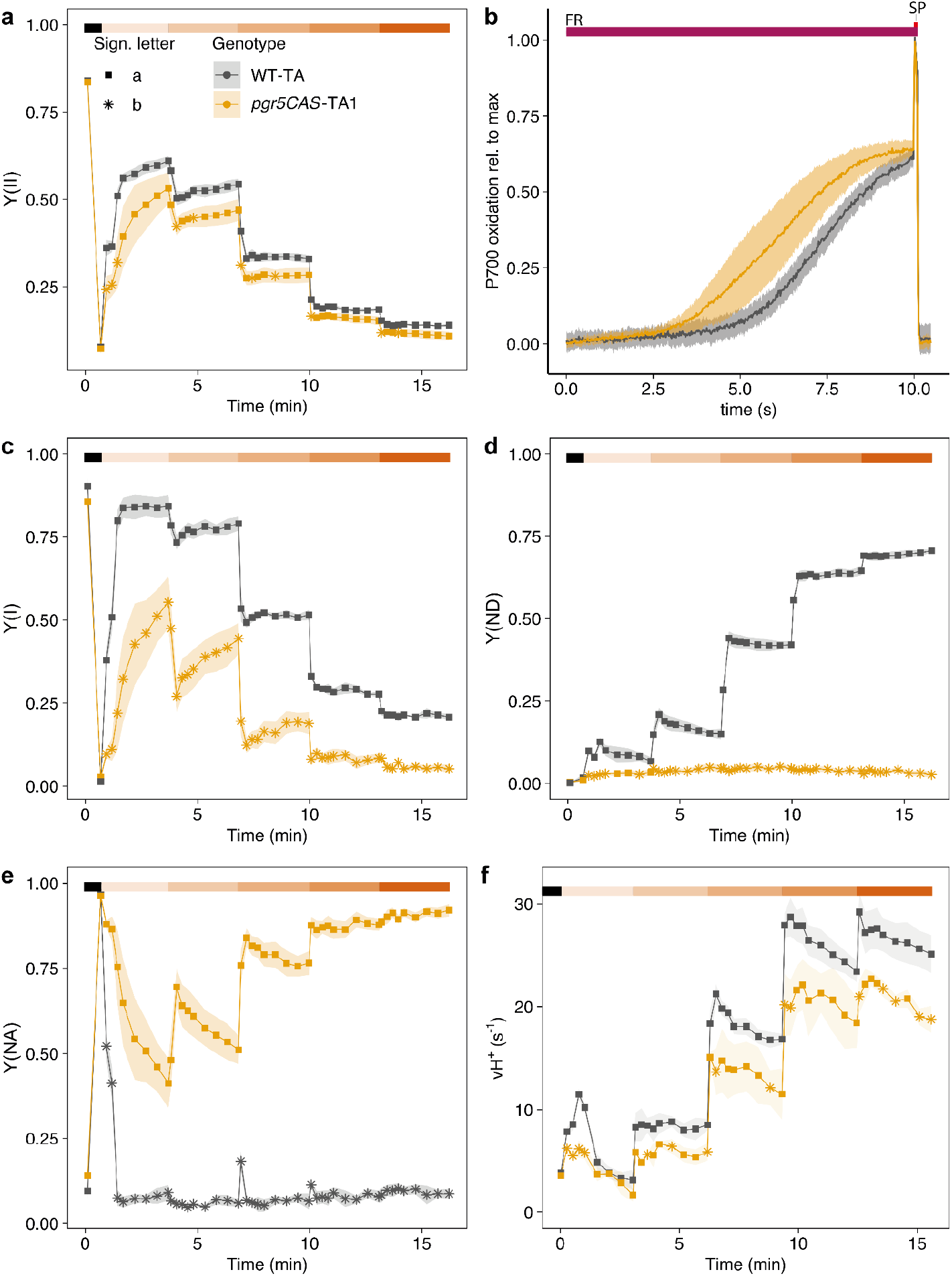
Photosynthetic parameters of WT TKTP-ATeam and *pgr5CAS*-TA1 in response to increasing light intensities. a) Yield of photosystem II (Y(II)). b) Oxidation of P700 under far red light normalised to a saturating flash at 10s. c) Yield of photosystem I (Y(I)). d) Donor-side limitation of PSI, also referred to as P700 oxidation (Y(ND)). e) Acceptor-side limitation of PSI, also referred to as P700 reduction (Y(NA)). f) Proton flux through the thylakoid membrane. Data points represent the mean of at least 3 biological replicates and shaded areas represent SEM. Different symbols indicate significant differences between treatments or genotypes at each time point calculated from a Tukey HSD test, alpha = 0.05. Coloured bars at the top of the panel indicate red actinic light intensities. The same light intensities as in Fig 2 a-d were used.

**Figure S4.**
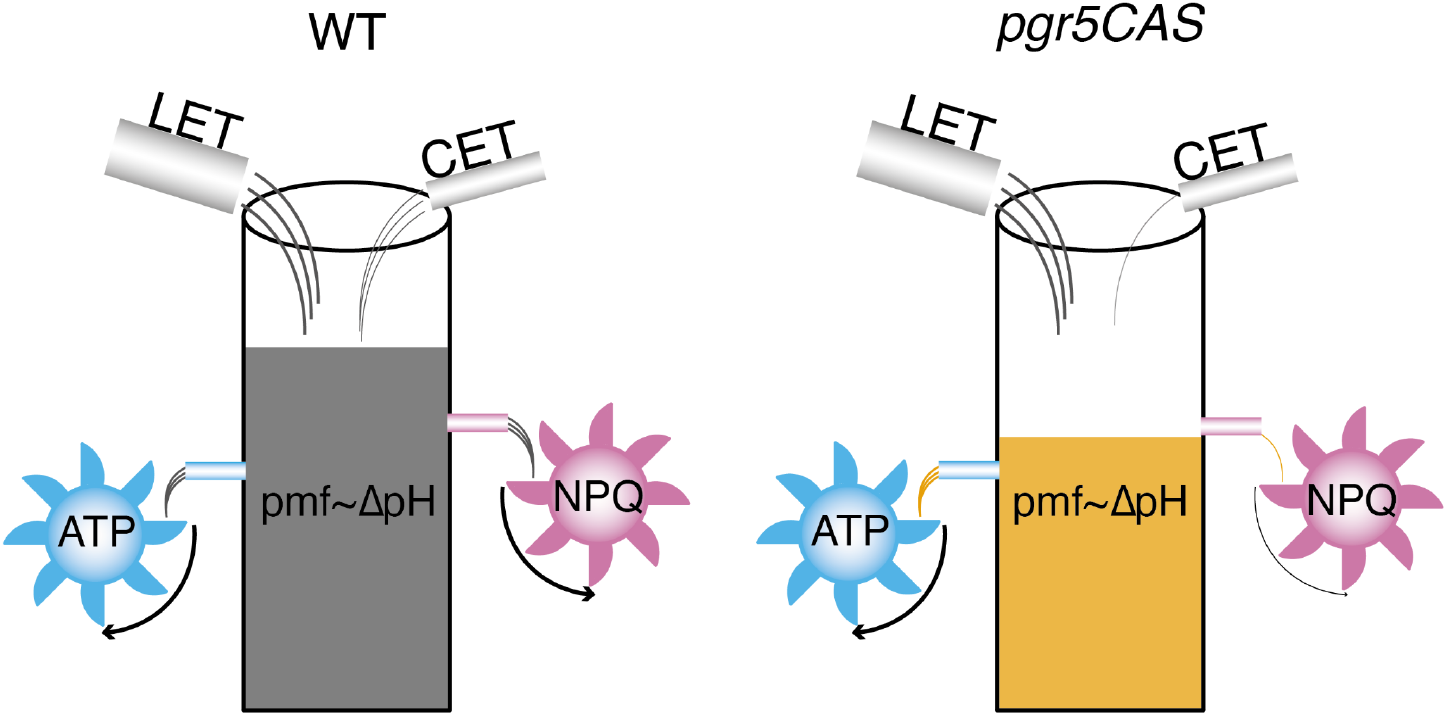
Clarified role of PGR5-dependent CET. In the WT, LET and CET drive the build-up of sufficient pmf for synthesis of ATP and induction of NPQ. In *pgr5CAS*, lack of CET results in a lower pmf, sufficient for ATP synthesis but not induction of significant NPQ, resulting in photoinhibition.

## Notes

### Competing Interest Statement

The authors have declared no competing interest.

